# Farmers’ awareness and occurrence of *Meloidogyne* species in tomato production in central Nepal

**DOI:** 10.1101/2025.08.01.667928

**Authors:** Jenish Nakarmi, K.C. Gopal Bahadur, Florian M.W. Grundler

## Abstract

Tomato (*Solanum lycopersicum*) is one of the most economically important vegetables in the world. Root-knot nematodes (RKN) form a complex of species that can cause severe losses in tomatoes. Since symptoms and damage depend on the particular nematode species, accurate species identification is critical for implementing potential control measures. Detailed surveys of plant parasitic nematodes in Nepal have not been conducted. Therefore, there is no information on which RKN species occur in key tomato-growing areas. We conducted an initial survey to assess the occurrence and importance of RKN. Nine different districts of Nepal were included: Bhaktapur, Chitwan, Dhading, Dolakha, Kaski, Kathmandu, Kavrepalanchok, Lalitpur, and Lamjung. In the first approach, 70 farmers were interviewed about their awareness and knowledge of RKN. More than 60% of the farmers surveyed knew of RKN and the main signs and symptoms. 40% had limited or no knowledge about it. In a second approach, we conducted a survey of RKN occurrence in tomato fields in the same nine districts. Soil sampling and subsequent analysis in the Kathmandu district revealed RKN infestation at very high levels. Except for Kaski and Chitwan, RKN prevalence was found to be 100% in the districts sampled. The overall prevalence of *Meloidogyne* species was also 100% in most districts. To identify the species, samples with RKN-infected galls were collected and determined morphologically using perineal patterns and molecularly using specific PCR analyses. Both methods showed that *Meloidogyne incognita* was the most abundant in the tomato fields, followed by *M. arenaria* and *M. javanica*. Our results confirm the importance of RKN in Nepal and suggest that it would be highly beneficial economically to increase farmer awareness of nematode problems and possible control measures.

## Introduction

Tomato (*Solanum lycopersicum* L.) is one of the major and economically important vegetable crops throughout the world (Quinet et al., 2019). It is a source of valuable nutrients, minerals, vitamins, and antioxidants (Bergougnoux, 2014). Global tomato production in 2019 was 181.89 million tons on 4.99 million ha with an average yield of 36.45 t/ha (*FAOSTAT*, 2021). According to national statistics, total tomato production in Nepal in the 2018/19 marketing year was 400000 tons on 23000 ha of land with a total yield of 17.39 t/ha in fiscal year (MoALD, 2020). The average consumption of tomatoes is 11.97 kg per person per year (Ghimire et al., 2017).

Of the 77 districts in Nepal, 22 are considered potential tomato-growing areas (Devkota et al., 2018). Open field production is common in sub-tropical Terai from November to March, while cultivation in mid and higher altitudes is done under plastic tunnels from April to September (Chaulagai & Koirala, 2021). Tomato cultivars are very sensitive to climatic and micro-climatic conditions, so adapted cultivars are used for different cropping systems and geographical areas (Chapagain et al., 2011). Production in plastic tunnels is based on improved hybrid plastic houses (Ghimire et al., 2001). Due to open borders and little regulation, many tomato varieties are in principle available to Nepalese farmers, but only four varieties are released by the National Seed Board in association with the Nepal Agricultural Research Council (NARC) (Magar & Gauchan, 2016).

Tomatoes are grown in Nepal under different socio-economic conditions. Many families produce tomatoes in subsistence agriculture; however, some farmers specialize in commercial tomato production. Local production competes in the market with imported products, which leads to limited profitability of domestic cultivation. In addition to small farm size, high pre-harvest and post-harvest losses also reduce the profitability of tomato production (Tiwari et al., 2020).

During cultivation and post-harvest storage, tomato plants are exposed to more than 200 diseases that can cause yield and quality losses. Most diseases affecting tomatoes in Nepal are caused by fungi, bacteria, viruses, and nematodes (Manandhar et al., 2020). RKN is one of the most devastating and widespread pathogens of tomato (Manandhar et al., 2020). The yield losses they cause depend on the nematode species, population density or infestation level, host plant, and factors such as plant nutrition and water supply (Ornat & Sorribas, 2008). In Nepal, data and information on RKN infestation and damage to crops are still very limited, for example, reports of *M. graminicola* occurrence in rice-wheat production and RKN in tomato crops at Hemja in Kaski district (Baidya et al., 2017; Pokharel et al., 2007). In addition to limited data, many farmers in Nepal lack awareness of PPN and information on control measures such as effective pesticides or technical devices. RKN are difficult to control once established in the field due to their wide host range and soil-borne nature (Baidya et al., 2017). Farmers who are unaware of the presence of RKN in their fields also do not consider preventive measures or management strategies. Therefore, the damage caused by RKN can affect crop yields and quality and lead to low income, food insecurity, and livelihood threats. To reduce severe crop losses and implement effective integrated management strategies, farmers need to be educated about RKN and nematode control and prevention.

The objective of this study was to survey farmers’ knowledge about the occurrence of RKN in their tomato crops and to analyse which *Meloidogyne* species are present in Nepalese tomato production. In this way, we attempt to fill existing information gaps and pave the way for improved management in tomato production in Nepal.

## Materials and methods

### Selection of tomato growing areas for the RKN survey and sampling

The RKN survey was conducted covering 14 various locations across 9 different districts (Bhaktapur, Chitwan, Dhading, Dolakha, Kaski, Kathmandu, Kavrepalanchok, Lalitpur, and Lamjung) in 2017-2018 (Figure 1). Among all, 2 are located in Terai region whereas rest all are located in Hilly region, covering diverse topographic regions of Nepal (Figure 2). These districts are all important tomato-growing areas in Nepal. The selected districts are located in different agro-ecological zones. In each district, 2-3 sites were selected for the survey and sampling.

**Figure 1:**
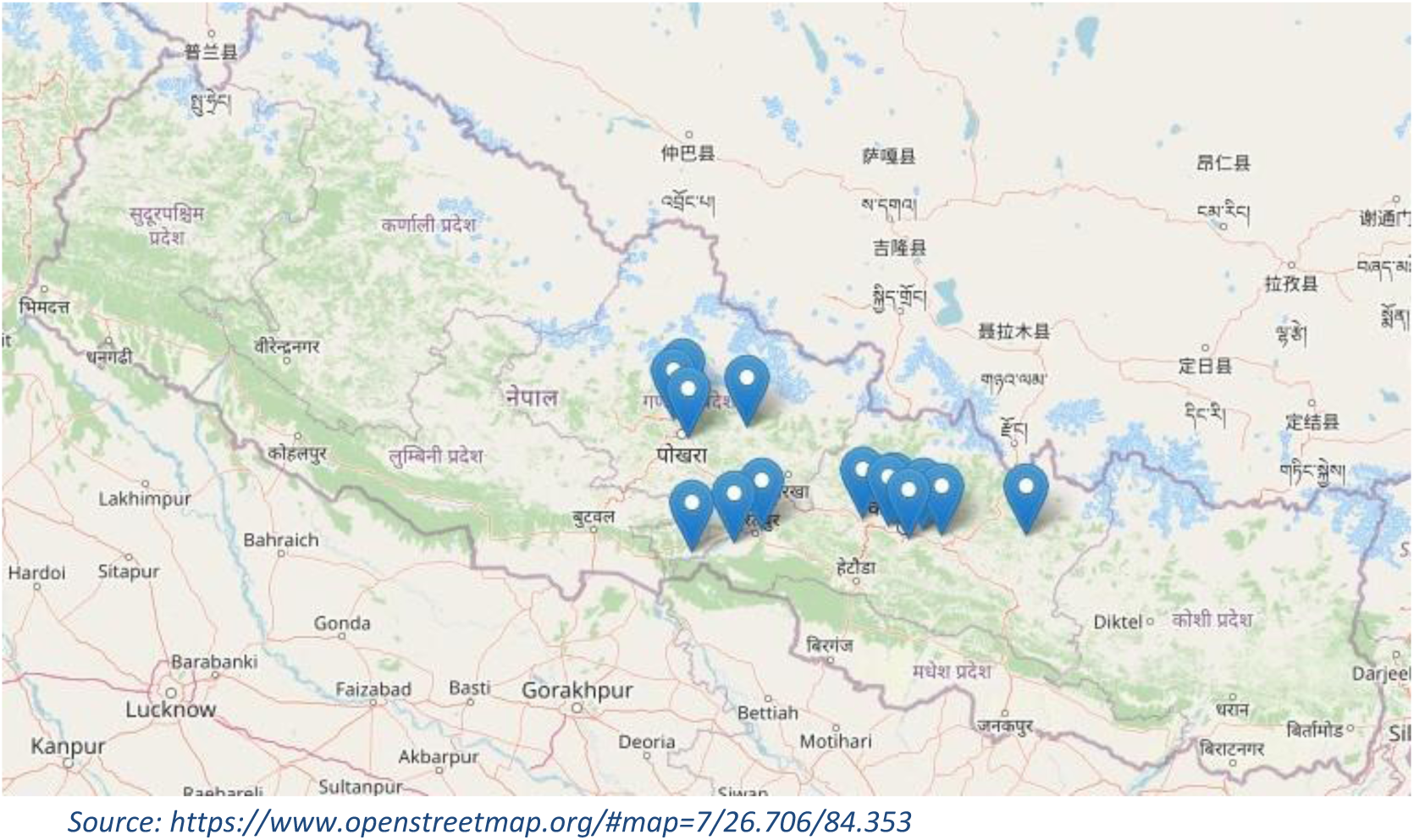
Map of Nepal indicating 14 locations where the survey was carried out and that are major tomato-growing areas.

**Figure 2:**
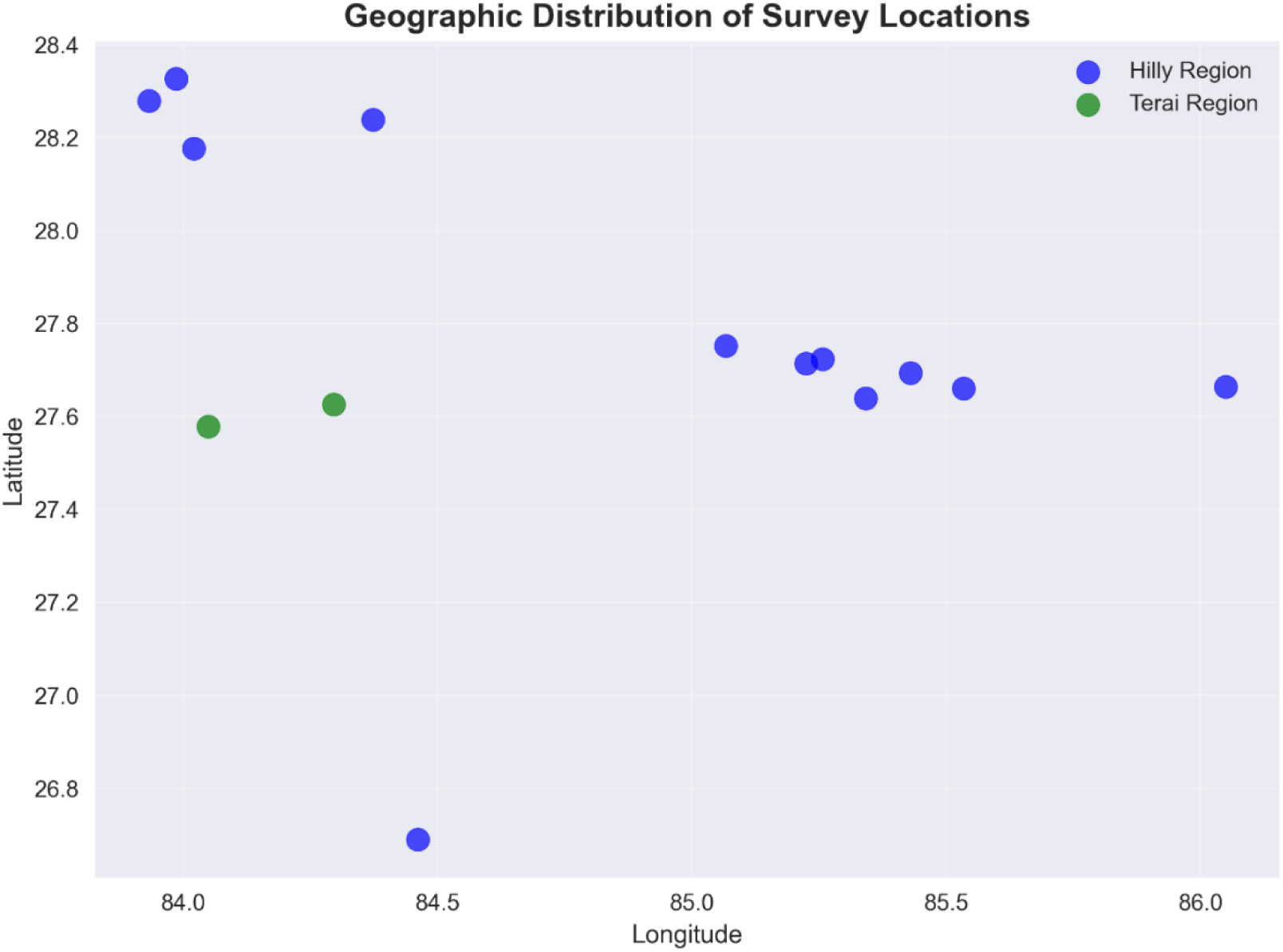
This scatter plot shows the geographical distribution of survey locations indicating different agro ecological zones of the respective location, based on latitude and longitude coordinates. Most locations (blue) are clustered between 27.7 and 28.3 latitude and 84.5 to 86.0 longitude, with fewer Terai locations (green) at lower latitudes (around 27.6 to 27.8).

Samples were collected from the 9 selected districts in open fields in the Terai and in plastic tunnels with an area of 75 m^2^ and a height of 2.5 m in the hilly and high-altitude regions of Nepal. 5 root samples infected with RKN were randomly collected from each field studied.

### Survey on farmers’ knowledge about RKN

70 farmers were interviewed using a structured questionnaire (Table 1). Face-to-face interviews were conducted to determine knowledge about RKN and the impact of RKN on tomato production. The questions were related: (i) the optimal season for tomato production (ii) different vegetables in the rotation (iii) tomato varieties (iv) knowledge of different diseases responsible for crop losses in tomatoes (v) knowledge about RKN in their field (vi) information on RKN symptoms, their description, severity, pathogenicity and host range (vii) management practices to control RKN (viii) evidence of biological control measures (ix) use and storage of chemical pesticides and (x) knowledge of the effects of pesticide on users, consumers, and the environment.

**Table 1:** Socio-demographic attributes of the growers with gender, age, educational status, and occupation.

| **Variables** | **No. of farmers** | **Percentage (%)** |
| --- | --- | --- |
| **Gender** |  |  |
| Male | 54 | 77 |
| Female | 16 | 23 |
| **Age (years)** |  |  |
| 16-25 | 0 | 0 |
| 26-35 | 15 | 21 |
| 36-45 | 28 | 40 |
| 46-55 | 23 | 33 |
| 55 and above | 4 | 6 |
| **Educational status** |  |  |
| Literate | 7 | 10 |
| Primary School | 0 | 0 |
| High School | 34 | 49 |
| Higher Secondary School | 29 | 41 |
| **Occupation** |  |  |
| Agriculture | 70 | 100 |
| Government employee | 0 | 0 |
| Business | 10 | 14 |
| Teaching | 0 | 0 |
| Retired | 10 | 14 |

No. of root samples (n) =70; no. of districts where samples were collected =9

### Analysis of the severity of RKN

Infested roots were placed in plastic bags, labelled, and taken to the Central Agricultural Laboratory (CAL) (Soil, Seed, and Crop Protection). To evaluate root galls, the entire root system of the collected plants was examined for the presence of galls. The root gall index (GI), based on the percentage of galled roots, and was used to evaluate infection. (Speijer & De Waele, 1997).

### Identification of *Meloidogyne* species

Morphological and molecular identification of *Meloidogyne spp.* was performed at the Laboratory of Molecular Phytomedicine, University of Bonn (Germany). Infected root samples were preserved in lactic acid (45%) for morphological analysis, and a similar number of infected root samples were preserved in absolute ethanol (99%) for DNA analysis.

### Morphological analyses by preparation of perineal patterns

Perineal patterns of females were used to identify RKN species as described by Taylor & Netscher, 1974. Female nematodes were collected from tomato roots preserved in lactic acid, and their posterior end was cut off with a fine and sharp blade. The body tissues were carefully removed, and the clean cuticle was transferred to a drop of glycerol, where it was carefully cut off. The posterior end of the females, including the vulva with the typical perineal pattern, was then transferred to a slide in a drop of glycerol. The mounted sections were covered with a coverslip and sealed with nail polish. The perineal pattern of the specimens was examined under the microscope (Leica) and compared with standard diagrams for species identification. Five perineal patterns from each specimen were examined for species identification.

### Molecular identification PCR analysis

To confirm species identification of *Meloidogyne* species, molecular analysis was performed on females collected from fields and greenhouse for comparison. Total genomic DNA was extracted from preserved females in ethanol. Ten females were used per root sample. A single female preserved in ethanol was immersed in 50 µl of sterile water. It was then crushed with a sterile toothpick. An aliquot of 2 µl of the suspension was used as a template for PCR reactions. To facilitate identification, a genomic DNA of standard and previously identified *M. incognita*, *M. javanica*, *M. arenaria,* and *M. graminicola* cultures were extracted from stock cultures and used as control. Identification of *Meloidogyne* species was based on the protocol for amplification of the mitochondrial NAD5 gene using primers NAD5F2 (TATTTTTTGTTTGAGATATATTAG) and NAD5RI (CGTGAATCTTGATTTTCCATTTTT) (Janssen et al., 2016). PCR was performed in a 25 µl reaction volume containing 16.25 μl of nuclease-free water (Sigma Aldrich ® Company), 5 µl of 5X green Gotaq buffer (Promega), 0.5 µl of dNTP mix (Promega), 0.5 µl of forward and reverse primer, 2 µl of template DNA and 0.25µl of Unit Taq DNA Polymerase (Promega). DNA amplification products were separated on a 1% agarose gel dissolved in 1×TAE and mixed with 5 ml PeqGreen (peclab) to 100 ml of agarose. Electrophoresis was performed at 80 volts for 60 minutes and visualized using UV light. The amplified DNA products were purified using NucleoSpin ^®^Gel and PCR Clean-up Kit (MACHEREY-NAGEEL) and quantified using a Nanodrop spectrophotometer. The purified PCR products were sequenced in both directions using NAD5F2 and NAD5RI primers (GATC Biotech, Germany). BLAST analysis was performed with the DNA sequences in the NCBI database.

### Data Analysis

All participants were active(Coyne et al., 2018)ly involved in the survey and provided complete responses to the questions asked. Data analysis was performed by using Python 3.8.

## Results

Our results showed that there are two optimum seasons for tomato production: (i) summer season for greenhouse cultivation in all hilly and high altitude districts, and (ii) winter season for open field production in Terai, i.e., Chitwan. It was found that 78.6% of farmers cultivated tomatoes in greenhouses in summer, while 21.4% of farmers cultivated them in open fields in winter (Figure 3A). However, the distribution of locations clearly showed a concentration of survey sites in the Hilly region and limited the coverage in Terai. In terms of tomato varieties, “Srijana” was the most popular; it was grown in plastic tunnels by the majority of farmers (79%), followed by “Samjhana” (24%), mostly in hilly regions. Farmers in Chitwan (22.9%) preferred to grow the variety “Surya-111” (Figure 3B), where tomatoes are produced in the open field.

**Figure 3:**
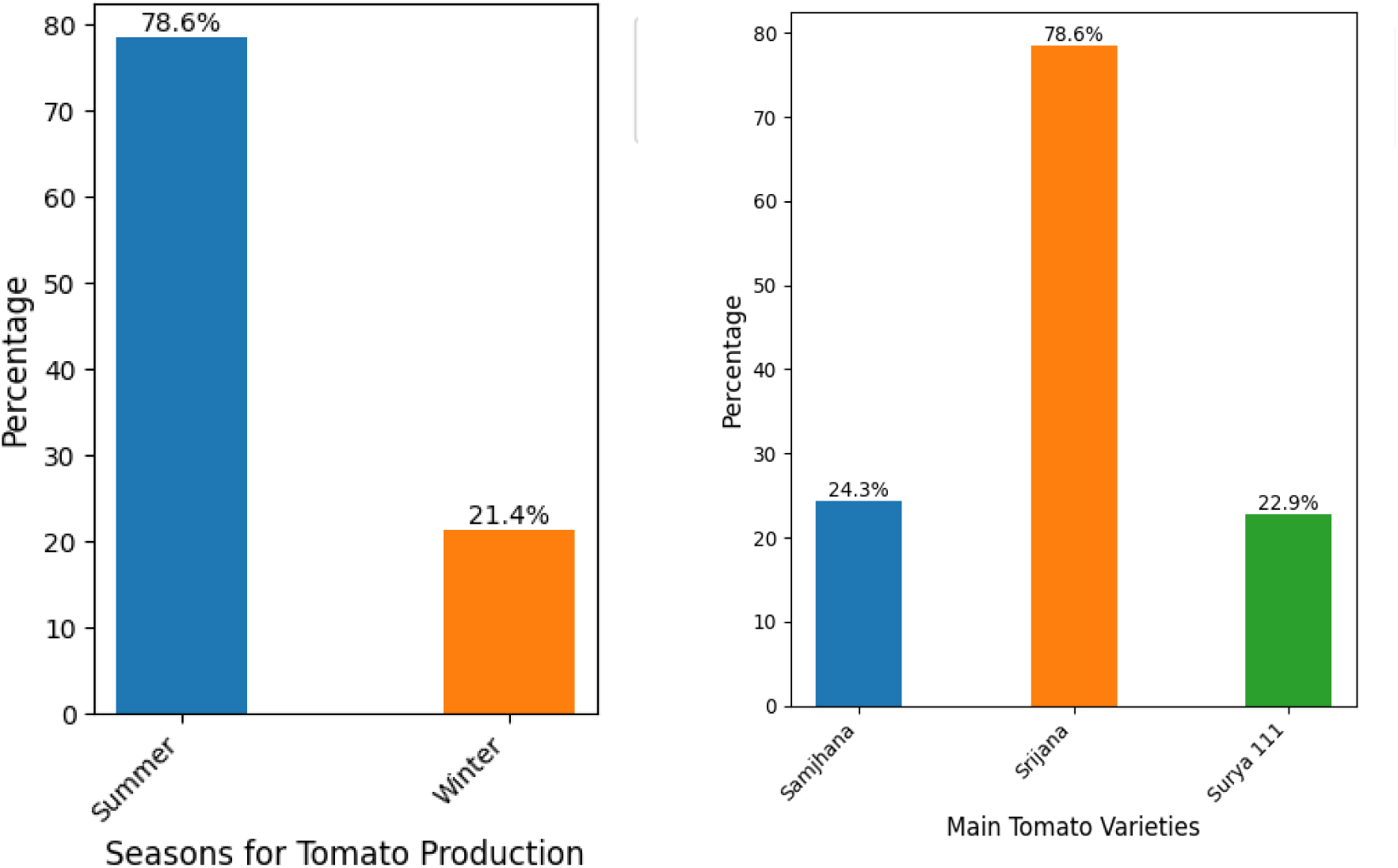
The graph shows the total percentage of interviewed farmers who produce tomato in two seasons, i.e. summer and winter. The graph shows the three tomato varieties cultivated by farmers in their fields

### Evaluation of the socio-demographic attributes of farmers

The socio-demographic characteristics of the 70 respondents with variables including gender, age, education, and occupation are shown in Table 1. The majority of farmers are men (77%), which shows a significant gender gap in agriculture. The involvement of women in tomato cultivation is low, reflecting limited access to farming resources and traditional gender roles. The interviewed farmers between 36 and 45 years group, is the largest of all, making up 40%. No farmers are aged 16-25, which indicates youth are not involved in farming, probably due to urban migration, education, or lack of interest in agriculture. Farming appears to be dominated by middle-aged adults, with relatively few older participants. A significant majority of the farmers (90%) have completed at least high school education. No one has only primary education, suggesting a relatively educated farming population. This could influence their openness to new technologies, modern farming methods, and market engagement. All respondents are involved in agriculture, but some also identify as retired or involved in business, likely as secondary occupations. No respondents are government employees or teachers, emphasizing agriculture as the sole or primary livelihood. All of the farmers interviewed engaged in commercial tomato farming but also used a small part of the yields for household consumption.

### Farmers’ recognition of RKN and its damage to tomato farms

In the survey, only 67% of farmers can recognize RKN as a disease affecting tomato plants, which is comparatively fewer than the other four diseases and pests (Figure 4). However, all growers (100%) were aware of the occurrence of late blight (LB), which is caused by *Phytophthora infestans* alongside bacterial and fungal wilt. Similarly, 82.9% of them were aware of the tomato leaf miner (TLM), *Tuta absoluta* (Meyrick) (Lepidoptera: Gelechiidae).

Among the 61% of farmers who recognize RKN as a disease in the previous graph, their understanding of its symptoms is fairly consistent. Less fruiting is the most commonly identified symptom (67%), followed closely by plant wilting and root galls, each recognized by 66% of farmers. Yellowing is the least recognized symptom, with 61% awareness (Figure 5).

**Figure 5:**
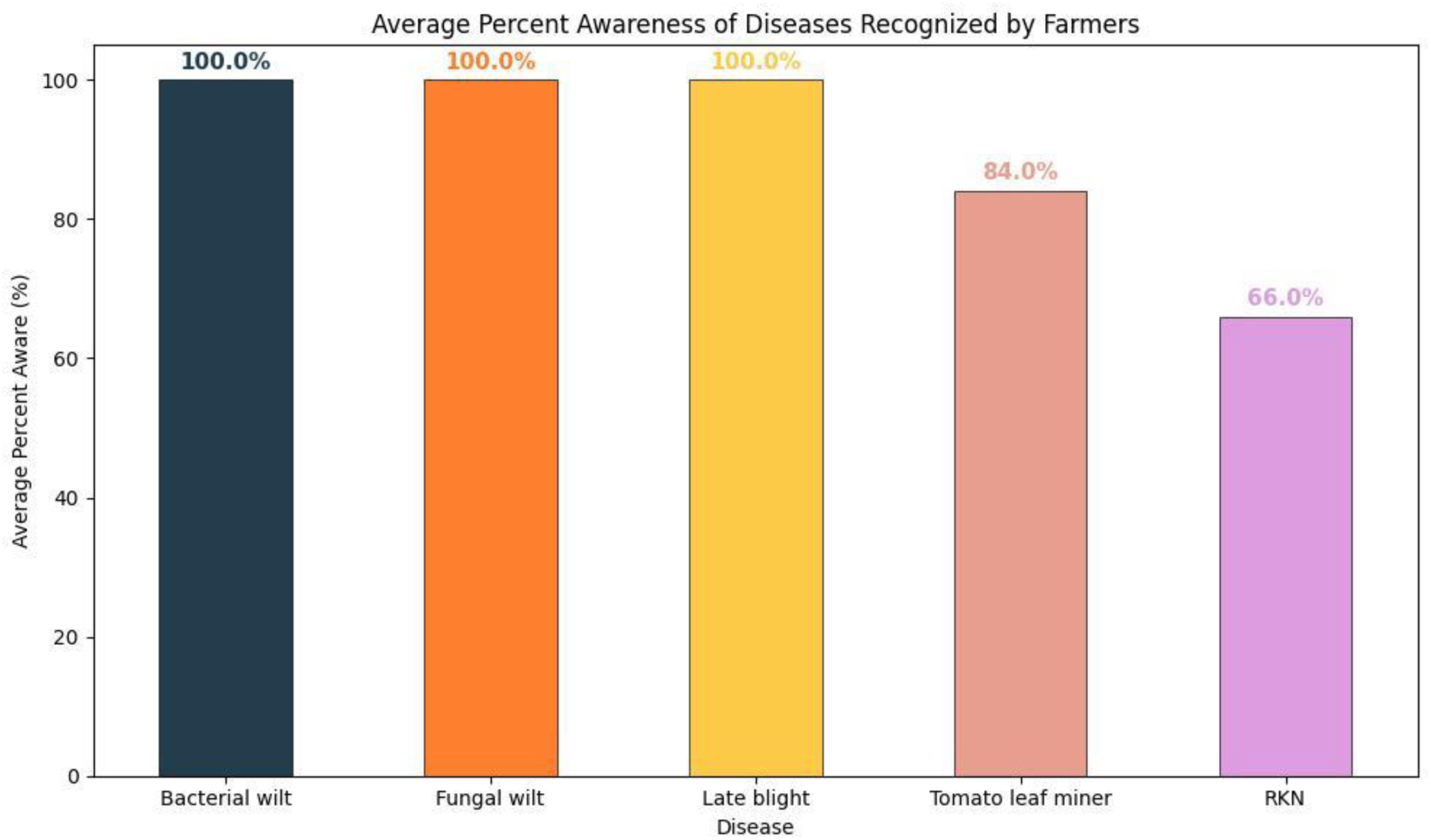
The graph shows the total percentage of interviewed farmers who recognized the diseases and pests that infected their tomato farming.

The farmers also reported experiencing yield losses in their tomato production. 33% of farmers were unaware of this impact. A majority of farmers (two-thirds) understand that RKN leads to reduced yields (Figure 6). However, a significant portion (one-third) still lacks awareness of the connection between RKN and production losses. The exact value of production loss was yet unclear.

**Figure 6:**
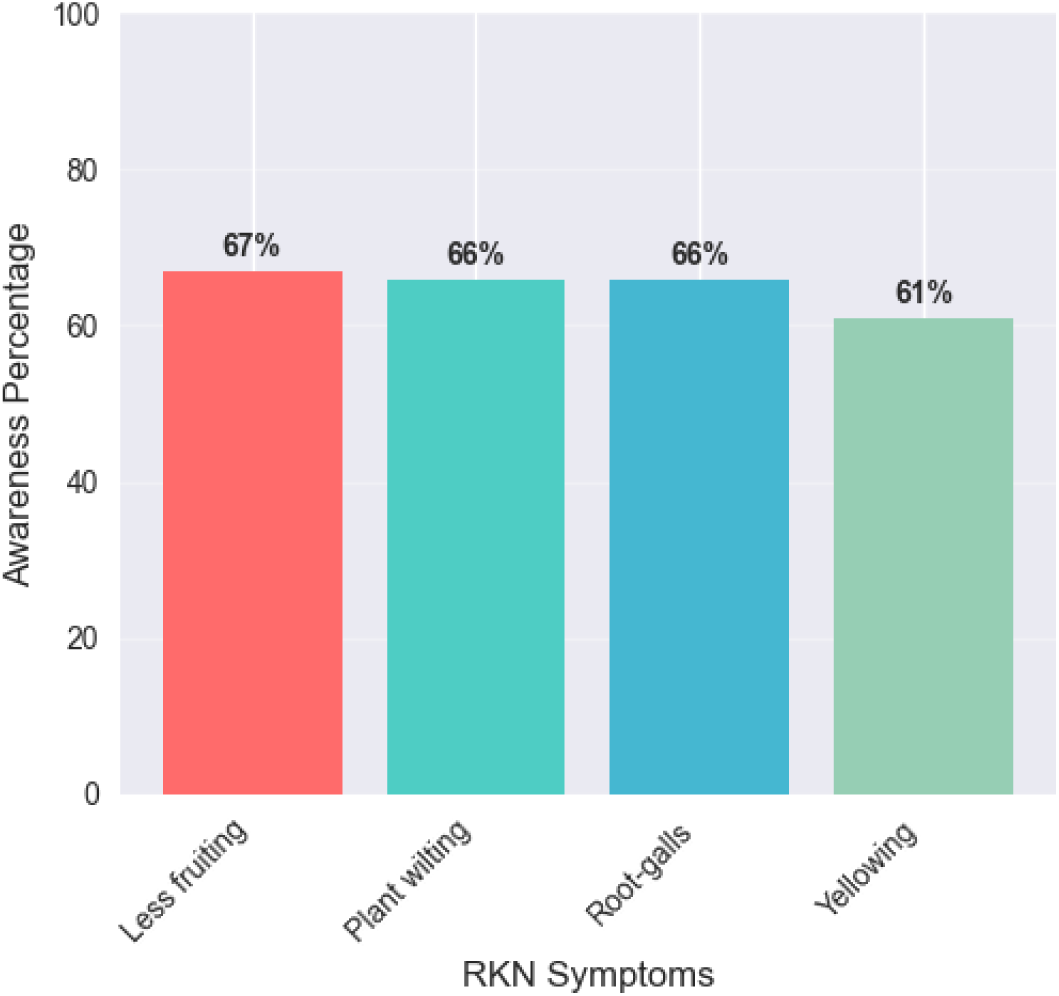
The graph shows the total percentage of interviewed farmers who were aware of RKN symptoms.

### Farmers’ knowledge on crops attacked by RKN

Regarding the vegetables grown in rotation, our survey found that cauliflower, coriander, black-eyed beans, string beans, and mustard greens were the most commonly grown (Figure 7). Cauliflower and coriander were most commonly grown by 56% of farmers, with cauliflower cultivation being most popular in Kathmandu and Kaski, while farmers in Bhaktapur grew more coriander. 46% of farmers surveyed in Chitwan grew black-eyed beans. String beans were among the most commonly grown vegetables in Kathmandu and Dhading (31% of farmers). 23% of farmers in Kaski and Kathmandu grew mustard greens.

**Figure 7:**
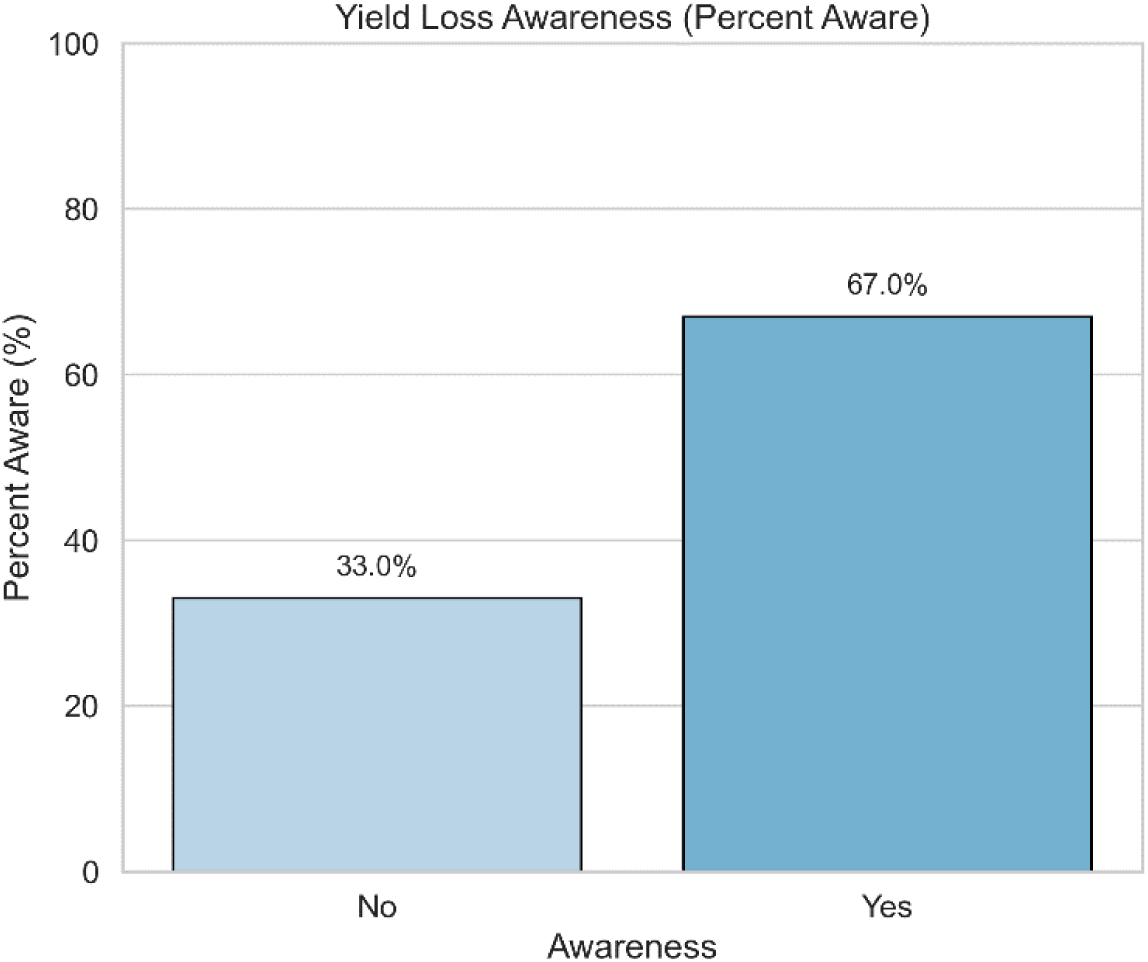
The graph shows the total percentage of interviewed farmers who encountered the yield loss in their tomato farms.

However, only 7% of respondents knew about alternative hosts, which refer to the variety of crops that RKN can infect (Figure 8). The awareness is very crucial for crop rotation planning against RKN.

**Figure 8:**
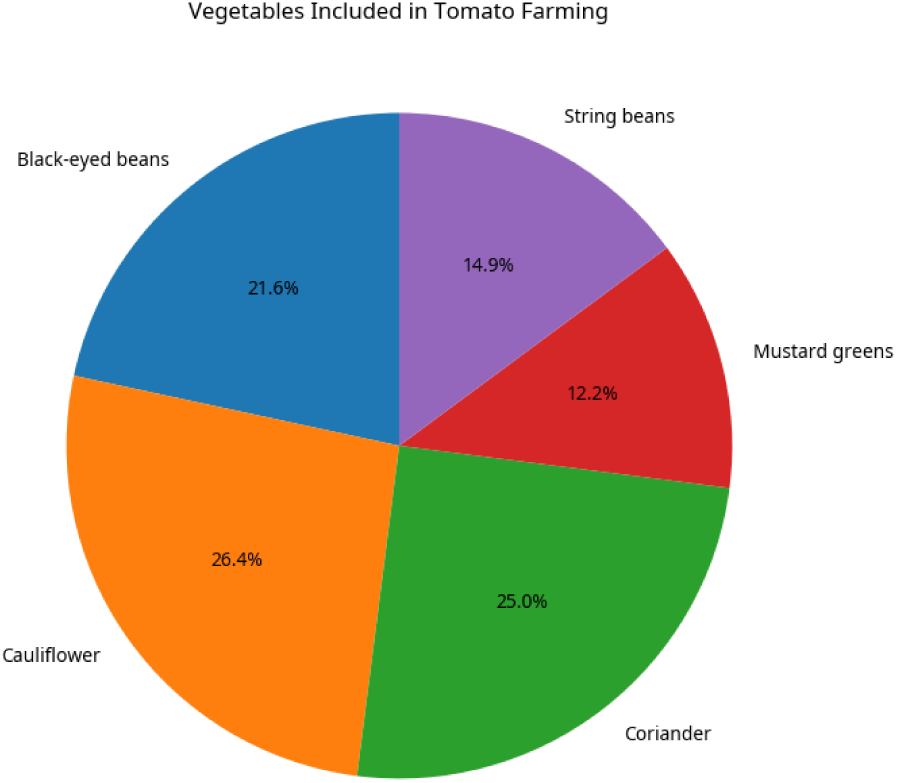
The pie-chart shows the total percentage of vegetables grown by farmers in their fields along or in rotation with tomato production.

### Management practices done by farmers against RKN suppression

Figure 9 presents a heatmap illustrating the adoption frequency (%) of different RKN management practices in response to key symptoms reported by farmers. The data reveal that chemical measures are consistently used by 57% of respondents across all symptoms, indicating their dominant role in current RKN control strategies. Names, recommended dosages, and details of the substances were not available as unknown chemical substances against RKN were used. In comparison, mustard cake (64%) and neem cake (61%) were also widely adopted, suggesting a growing preference for organic soil amendments. Mustard as a rotation or trap crop was used by 29% of farmers, representing a moderate level of awareness. However, marigold was the least adopted strategy, with only 7% of farmers using it regardless of symptom type. The uniformity in adoption rates across symptoms indicates a general treatment-based approach rather than symptom-specific interventions. These findings underscore the need for more nuanced extension services that promote integrated and symptom-targeted RKN management, especially emphasizing underutilized but ecologically beneficial practices like marigold intercropping.

**Figure 9:**
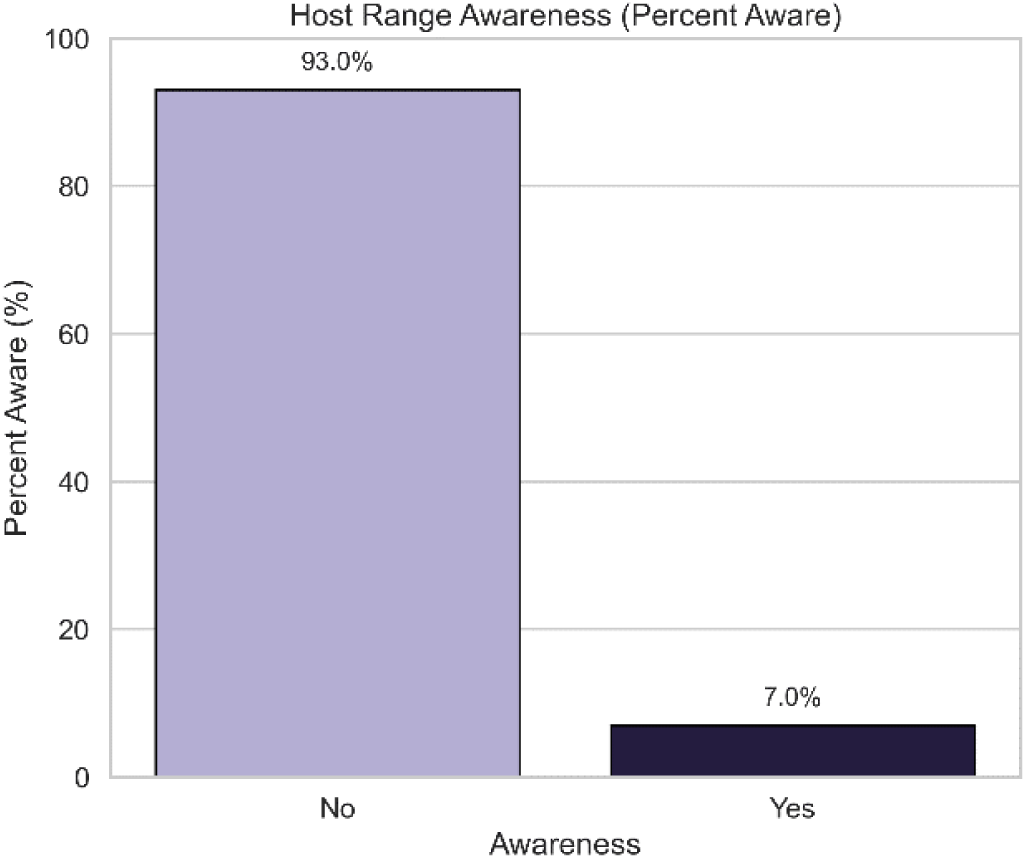
The graph shows the total percentage of interviewed farmers who have knowledge about the host range of RKN besides tomato.

In addition, farmers were asked for information on the potential negative impacts of chemical use (Figure 10). 20% of the farmers were aware of the possible hazardous effects of chemicals on the environment. None of the farmers were aware of the possible harm of killing non-target organisms from chemicals, but a total of 58% of farmers were aware of possible side effects on human health, with regional differences. In contrast, no farmer was aware of the existence of specific antagonistic microorganisms, such as bacteria or fungi that can be used to control RKN.

### Understanding the tomato farmers’ perception and responses to RKN infestation

The correlation matrix illustrates the interrelationships between farmers’ perceptions of RKN symptoms and their awareness and adoption of various management strategies Figure 11. Strong positive correlations were observed among key symptom variables such as root galls, plant wilting, yellowing, and less fruiting. This indicates that these symptoms frequently co-occur in the field. Among management strategies, the use of chemical measures exhibited moderate positive correlations with both neem cake and mustard cake, suggesting that farmers often apply multiple control methods simultaneously. Interestingly, awareness of environmental pollution and human health showed weaker correlations with direct management practices, potentially reflecting a gap in understanding of the ecological and health-related consequences of chemical usage. The low or negative correlation between host range and other variables further suggests limited farmer awareness of RKN host specificity.

**Figure 11:**
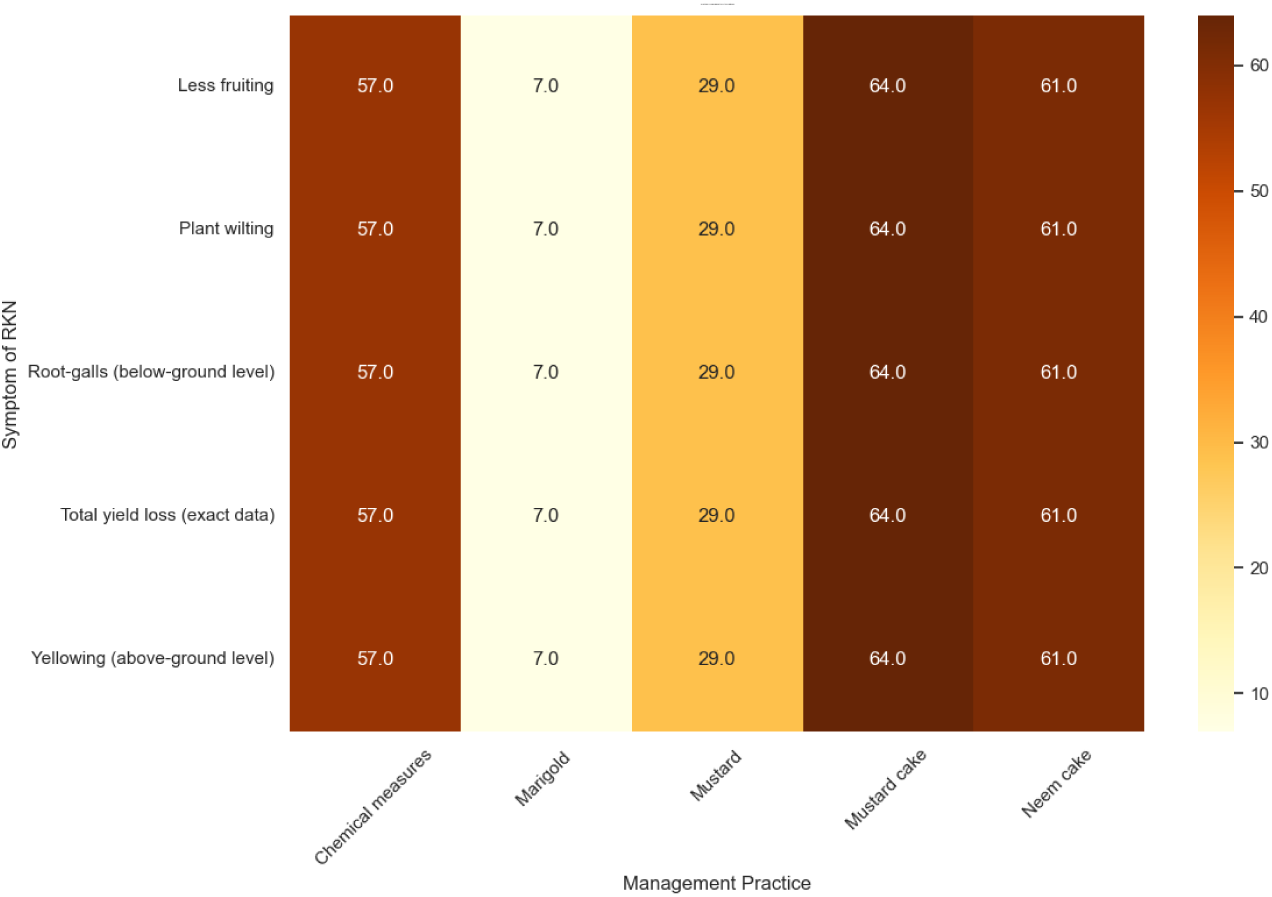
The heatmap visualizes the relationship between observed symptoms of RKN and the management practices adopted by farmers. It appears to represent the number or percentage of farmers who reported using each practice when a specific RKN symptom was observed.

### Severity of RKN

The severity of RKN infestation was assessed by the formation of root-galls by RKN, which were scored according to GI introduced by Speijer & De Waele (1997). To evaluate the GI, five root samples from each of the visited tomato fields were examined. The mean value of the observed severity was calculated from all the observed tomato plants in the surveyed districts (Table 2). The highest infestation level was observed in Kathmandu with 4, followed by Lalitpur with 4. The lowest infestation level was observed in Bhaktapur with 3.

**Table 2:** Incidence, prevalence, and galling index of root-knot nematodes in different sampling districts.

| Districts | Mean Galling index |
| --- | --- |
| Bhaktapur | 3 |
| Chitwan | 4 |
| Dhading | 4 |
| Dolakha | 4 |
| Kaski | 3 |
| Kathmandu | 4 |
| Kavrepalanchok | 3 |
| Lalitpur | 4 |
| Lamjung | 3 |

No. of root samples (n) =65; no. of districts where samples were collected =9

### Identification of the different *Meloidogyne* species

#### Morphological identification

For preliminary species identification, perineal patterns of female RKN were examined from 65 collected root samples from 9 districts. Some specimens could not be identified using perineal patterns. The perineal patterns of females of *Meloidogyne* were high, trapezoidal dorsal arch, and narrow dorsal curved (Figure 12a). Each five specimens was collected from Chitwan, Dolakha, Kathmandu, Kaski, Lalitpur, and Lamjung. These specimens were matched typically to *M. incognita* (Aydinli & Mennan, 2016). *M. incognita* is the most common species that infects tomato crops worldwide. Similarly, the specimens collected from Dhading were closer to *M. javanica* (Eisenback et al., 1980) (Figure 12b). These specimens included common lateral lines, which divide the dorsal and ventral marks. The perineal patterns of females collected from Bhaktapur and Kavrepalanchok had a low dorsal arch with forming shoulders. Lateral lines were distinct, dorsal and ventral striae connected with an angle and forked (Figure 12c). The dissected specimens of *Meloidogyne* were attributed to *M. arenaria* (de Araújo Filho et al., 2016).

**Figure 12:**
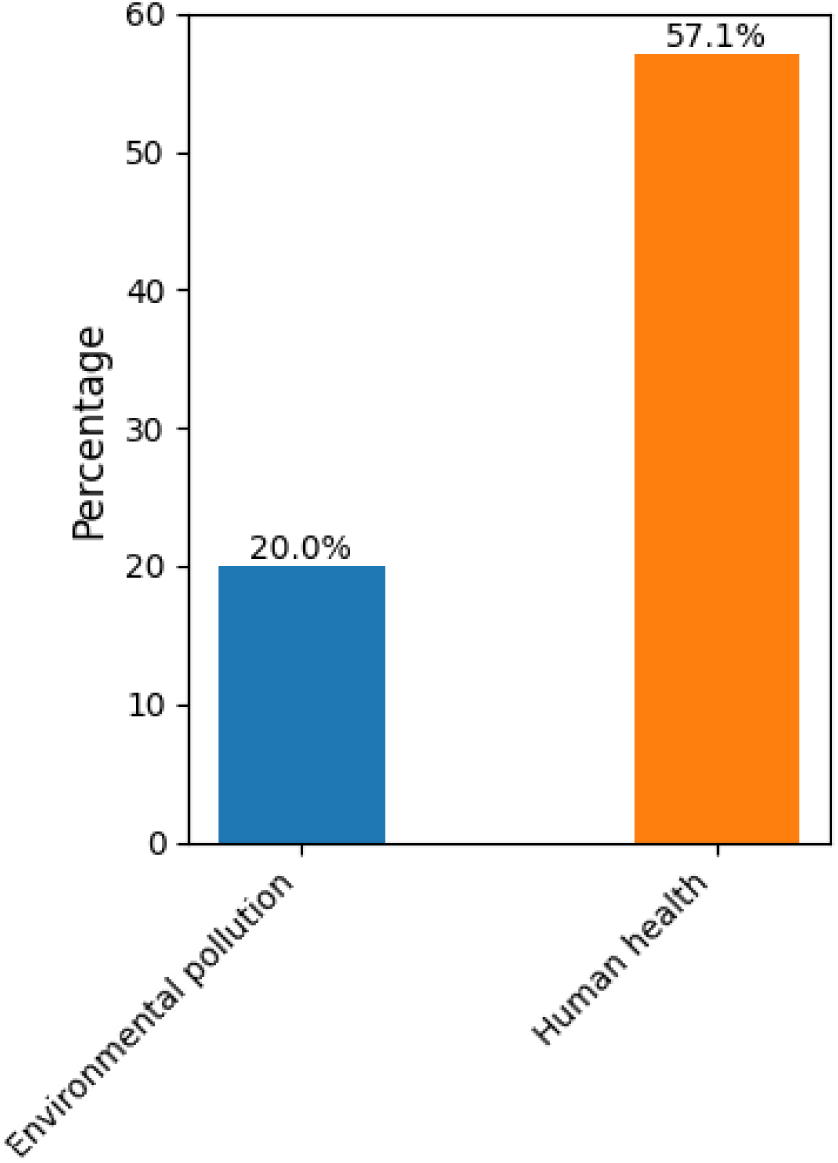
The graph shows the total percentage of interviewed farmers showing the knowledge on the adverse effects of chemicals used in suppression of RKN in their tomato farms.

**Figure 13:**
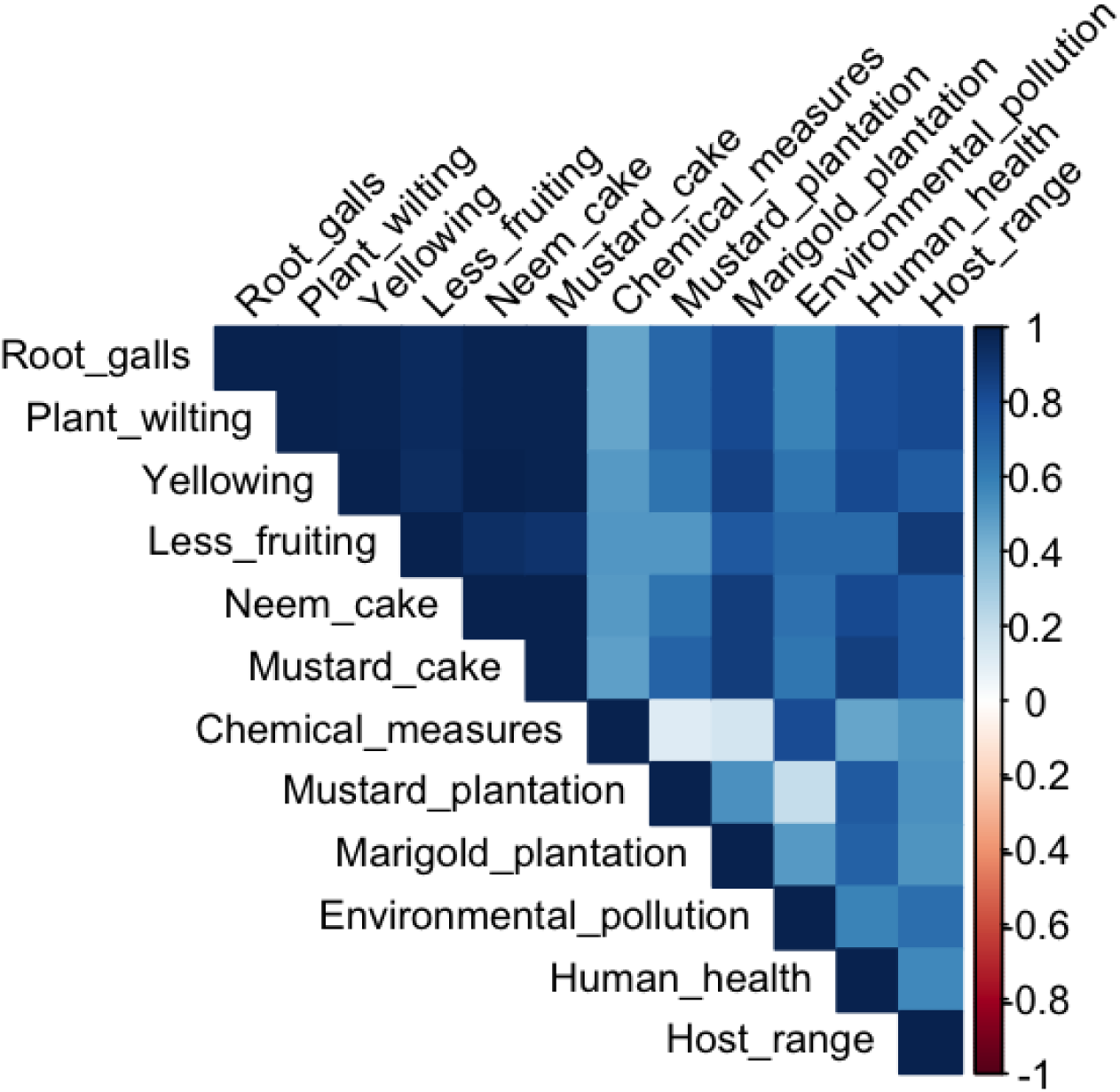
Correlation matrix showing relationships among RKN symptoms, management strategies, and awareness indicators. Positive correlations are shown in blue and negative correlations in red, with intensity reflecting the strength of correlation (Pearson’s r). Strong positive associations were observed between RKN symptoms (e.g., root galls, plant wilting, yellowing) and chemical measures, while awareness indicators such as human health and environmental pollution showed weaker or negative correlations with symptom-based responses. This suggests a non-targeted management approach and limited ecological awareness among farmers.

**Figure 14:**
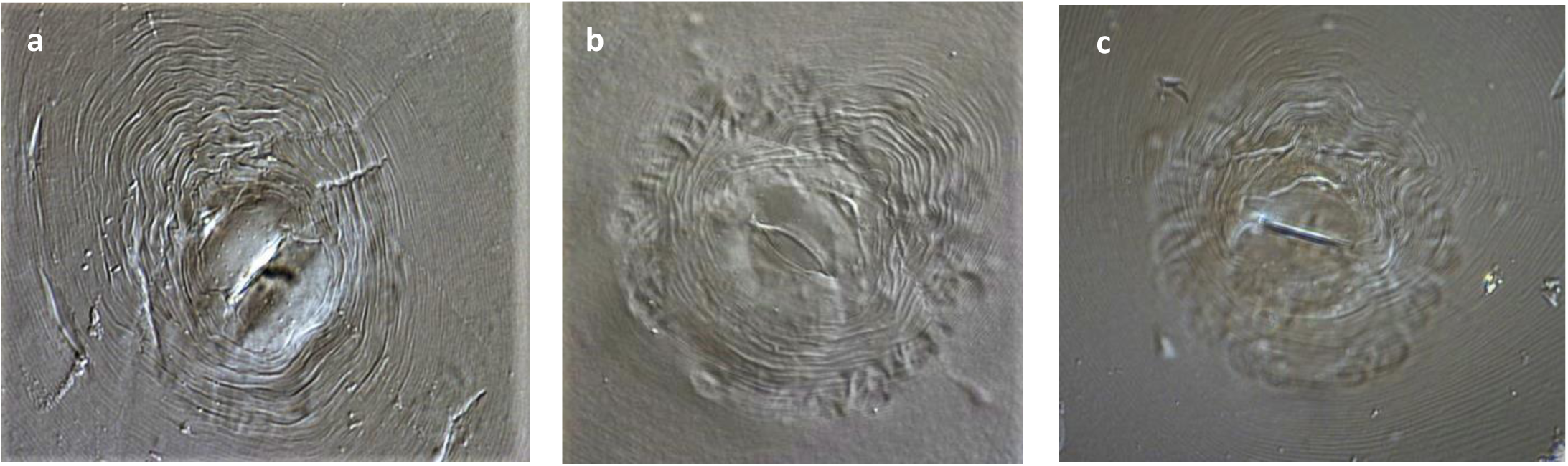
Photomicrographs of perineal patterns of RKN female: (a) *M. incognita* (b) *M. javanica* (c) *M. arenaria*.

#### Molecular identification

To confirm the morphological characterization, DNA sequence blasting and sequence of NAD5 gene were used to identify RKN species observed on the female perineal specimens. The identified NAD5 gene fragment was a reliable DNA marker for the identification of the most common tropical *Meloidogyne* species, i.e. *M. incognita*, *M. javanica* and *M. arenaria.* BLAST analysis revealed 100% identity with the sequence of *M. incognita*, *M. javanica,* and *M. arenaria,* respectively. *M. incognita* was predominantly detected at the sample locations: Chitwan, Dolakha, Kaski, Kathmandu, Lalitpur, and Lamjung (Table 3). The results of the BLAST analysis also confirmed the female perineal patterns observed in the following districts. Similarly, *M. arenaria* was detected in the samples from Bhaktapur and Kavrepalanchok districts (Table 3). The female perineal patterns observed in both districts also confirmed the same species. *M. javanica* was detected only in Dhading district (Table 3), which also confirmed the results of the morphological characterization of the female samples.

**Table 3:** Sampling districts with locations, agro-ecological zones and identified *Meloidogyne spp*.

| Districts | Locations | Identified *Meloidogyne spp.* |
| --- | --- | --- |
| Bhaktapur | Jhaukhel | *M. arenaria* |
| Bhaktapur | Kharipati | *M. arenaria* |
| Chitwan | Chanauli | *M. incognita* |
| Chitwan | Gunjannagar | *M. incognita* |
| Dhading | Mahadevbesi | *M. javanica* |
| Dolakha | Charikot | *M. incognita* |
| Kaski | Puranchaur | *M. incognita* |
| Kaski | Hemja | *M. incognita* |
| Kaski | Chhinedada | *M. incognita* |
| Kathmandu | Dahachok | *M. incognita* |
| Kathmandu | Ramkot | *M. incognita* |
| Kavrepalanchok | Nala | *M. arenaria* |
| Lalitpur | Harisiddhi | *M. incognita* |
| Lamjung | Beshisahar | *M. incognita* |

No. of root samples (n) =65; no. of districts where samples were collected =9

## Discussion

### Sampling limitations and socio-demographic trends in tomato production

The survey locations and samples of our study may be biased to some extent due to limited contacts, problematic infrastructure, and the current political instability in the country (Devkota et al., 2018; Pokharel & Acharya, 2015). These constraints likely influenced the geographical distribution of respondents, with a predominance of the Hilly regions and limited coverage in Terai, which may affect the generalizability of findings.

The predominance of middle-aged farmers (77%) highlights a significant gender imbalance and generational gap in agriculture, consistent with trends in Nepal and South Asia, where women often face restricted access to land and decision-making despite their crucial roles (FAO, 2011). With no representation from youth in commercial tomato farming may mirror broader national patterns of rural-urban migration and declining youth interest in farming (Lutuf et al., 2018; Phadera, 2016; Thapaliya et al., 2023). Encouragingly, the high educational attainment (90% with at least secondary-level education) suggests a potential for effective knowledge transfer if appropriate training is provided.

### Farmers’ knowledge and awareness of RKN

The survey revealed that only 67% of farmers were able to recognize RKN as a harmful pest affecting tomato crops. This recognition rate is considerably lower than for other common diseases like LB and TLM, causing highly visible damage (Ye et al., 2015). The nematode awareness remains still limited compared to fungal, bacterial and pests threats. Similar findings have been reported in other parts of South Asia, where RKN is often under-recognized due to its below-ground impact and gradual symptom development (Coyne et al., 2018; Luc et al., 2005).

Although 67% could identify RKN, symptoms were less recognized. This indicates a basic but incomplete understanding of the symptom complex of RKN, limiting the ability of farmers to take early and accurate action. Furthermore, about one-third of farmers did not link RKN with yield losses, highlighting a critical gap in awareness of its economic impact (Z. Khan et al., 2012).

### Understanding of RKN Host Range and Crop Rotation

A mere 7% of surveyed farmers were aware of the host range of RKN, a major knowledge gap that can undermine effective crop rotation strategies. This is especially concerning given that *Meloidogyne* spp. infect a broad spectrum of vegetable crops including solanaceous crops, legumes, brassicas, and umbel lifers such as cauliflower, beans, mustard, and coriander (Luc et al., 2005). Without a clear understanding of these alternative hosts, farmers may unintentionally facilitate nematode survival by rotating crops with similarly susceptible species (Jones et al., 2013). Research underscores that ineffective crop sequencing is a key contributor to persistent RKN infestations, whereas strategic rotations, such as okra–cowpea–cabbage, have shown success in suppressing nematode populations under field conditions (Khan et al., 2023). Conversely, repetitive cultivation of RKN-susceptible crops can perpetuate the nematode life cycle, leading to chronic root damage and declining yields (Jones et al., 2013). To address this, agricultural extension services must emphasize farmer education regarding RKN host specificity. Promoting the use of non-host crops or suppressive species such as marigold (*Tagetes spp.*) and cereals can reduce nematode populations naturally, offering a sustainable and cost-effective alternative to chemical nematicides (Luc et al., 2005). Educational outreach and farmer field schools should incorporate visual aids, rotation planning, and demonstrations to improve farmer understanding and adoption of nematode-suppressive practices.

### Knowledge gaps and trends in nematode management practices

The heatmap analysis revealed that chemical nematicides were the predominant management strategy, regardless of the specific symptom type. This indicates a non-specific, blanket application approach, likely stemming from limited awareness of nematode biology and integrated pest management (IPM) principles. Most farmers could not name the chemical products used nor cite appropriate application rates or safety precautions, underscoring a critical knowledge gap in safe pesticide use. Such uninformed practices carry serious risks, including the development of resistance, contamination of soil and water resources, and harm to human and environmental health (Luc et al., 2005; Sikora et al., 2018).

Encouragingly, the survey showed relatively high adoption of organic soil amendments, with mustard cake and neem cake frequently applied. These amendments are known for their nematostatic properties and soil health benefits (Baheti et al., 2019; Javed et al., 2008; Singh et al., 1996). However, their widespread use appeared more intuitive than strategic, as farmers uniformly applied all treatments regardless of the specific symptoms presented. This reactive approach reflects a lack of diagnostic capacity and a failure to select evidence-based treatments.

In contrast, only 7% of respondents practiced marigold (Tagetes spp.) intercropping, a well-established biological control method for root-knot nematodes (Mandal & Hossain, 2017). This extremely low adoption rate highlights poor dissemination of agro-ecological practices, despite their effectiveness in both field and protected cultivation systems. Moreover, solarization, another cost-effective and environmentally friendly technique, was not used by any farmer under plastic tunnel conditions, a missed opportunity for sustainable pest suppression (Sikandar et al., 2020).

Furthermore, only 20% of farmers recognized the environmental risks associated with chemical use, and none were aware of their impacts on non-target organisms, including beneficial nematode antagonists and soil microbes. While 58% acknowledged health risks to humans, regional differences suggest inconsistent access to training and advisory services. These patterns are consistent with other developing country contexts, where awareness is often limited to immediate health threats rather than broader ecological consequences (FAO, 2011).

Alarmingly, none of the farmers had any knowledge of biological control agents, such as *Paecilomyces lilacinus*, *Pochonia chlamydosporia*, or *Bacillus subtilis*, despite their proven efficacy in managing *Meloidogyne* spp. under diverse field conditions (Whipps & Davies, 2000).The complete absence of such knowledge represents a critical extension and education gap, particularly in light of the growing need for non-chemical, sustainable nematode control options.

Collectively, these findings stress the urgent need to reform extension services, shifting from product promotion toward knowledge-driven, IPM-based support systems. Capacity building must emphasize symptom recognition, ecological principles, and the integration of biological, cultural, and chemical control strategies (Hooks et al., 2010; Viaene et al., 2006). Future interventions should also include hands-on training, farmer field schools, and localized demonstration plots to increase adoption of effective, site-specific nematode management strategies.

### Severity of RKN Infestation

Root-knot nematode infestation levels were assessed using the Gall Index (GI), revealing moderate to severe infestations across most surveyed districts. These findings are consistent with economic threshold levels that have been linked to yield losses exceeding 50–60% in tomato production (Seid et al., 2015). The widespread presence of RKN across diverse agro-ecological zones reflects both their adaptability and the consequences of poor nematode management, particularly in high-value, year-round vegetable production systems. Contributing factors include monoculture practices, limited crop rotation, and indiscriminate chemical use, all of which favor nematode buildup and persistence in soil (Viaene et al., 2024). Moreover, the uniform severity across districts signals an urgent need for national-level nematode surveillance and integrated intervention programs, particularly in high-production regions like Kathmandu Valley. Without targeted management and diagnostics, RKN infestations are likely to intensify, jeopardizing both crop productivity and soil health in the long term.

### Identification and Distribution of *Meloidogyne* Species

This study represents the first documented effort in Nepal to integrate morphological diagnostics (perineal pattern analysis) with molecular techniques (NAD5 mitochondrial gene sequencing) for the identification of *Meloidogyne* spp. Three root-knot nematode (RKN) species were identified: *M. incognita*, *M. javanica*, and *M. arenaria*. Among these, *M. incognita* emerged as the most widespread and dominant, occurring in five of the seven surveyed districts: Chitwan, Kathmandu, Lalitpur, Kaski, and Dolakha. This distribution aligns with global patterns, where *M. incognita* predominates in solanaceous cropping systems, particularly in tropical and subtropical regions (Aydinli & Mennan, 2016; Eisenback et al., 1980; Jones et al., 2013).

*M. javanica* was identified exclusively in Dhading, and *M. arenaria* in Bhaktapur and Kavrepalanchok, suggesting localized ecological niches or crop rotation practices that may influence species distribution. The use of NAD5 gene-based molecular markers proved effective in confirming morphological diagnoses, supporting prior findings on their diagnostic robustness (de Araújo Filho et al., 2016).

Species-level identification is of critical management relevance, as *Meloidogyne* species differ significantly in host range, virulence, temperature tolerance, and interaction with host resistance genes (Hallmann & Kiewnick, 2018; Trudgill, 1997). For instance, resistance genes like *Mi-1.2* in tomato offer protection against *M. incognita* and *M. javanica*, but not against *M. enterolobii* or under high soil temperatures exceeding 28 °C (El-Sappah et al., 2019; Przybylska & Obrępalska-Stęplowska, 2020). This highlights the need for species-specific management strategies, particularly as climate change may influence nematode population dynamics and species composition.

## Conclusion

The findings of this study highlight serious gaps in farmers’ knowledge of RKN biology, host range, and sustainable management practices. While some organic methods are in use, a lack of understanding about the pest’s lifecycle, alternative hosts, and safe control options undermines long-term suppression efforts. Training farmers in symptom recognition, biological control, and IPM practices particularly tailored to species-specific nematode threats will be crucial for minimizing yield losses and promoting ecological sustainability in Nepal’s commercial tomato farming sector.

## Acknowledgments

The authors would like to thank the Institute of Agriculture and Animal Sciences (IAAS), Tribhuwan University (Lalitpur, Nepal), Mr. Shiva Yendo and Mr. Khem Paudel for the additional support in surveying Nepal, and all the farmers for their kindness and trust. The authors also like to thank Dr. Oliver Chitambo and Dr. Samer Habash for their valuable laboratory help.

## References

Aydinli, G., & Mennan, S. (2016). Identification of root-knot nematodes (Meloidogyne spp.) from greenhouses in the Middle Black Sea Region of Turkey. Turkish Journal of Zoology, 40(5), 675–685. 10.3906/zoo-1508-19

Baheti, B. L., Bhati, S. S., & Singh, H. (2019). Efficacy of Different Oil-Cakes as Soil Amendment for the Management of Root-knot Nematode, Meloidogyne incognita Infecting Okra (Abelmoschus esculentus L.). International Journal of Current Microbiology and Applied Sciences, 8(12), 799–808. 10.20546/ijcmas.2019.812.104

Baidya, S., Timila, R. D., KC R. B., Manandhar, H. K., & Manandhar, C. (2017). Management of root knot nematode on tomato through grafting root stock of Solanum sisymbriifolium. Journal of Nepal Agricultural Research Council, 3, 27–31. 10.3126/jnarc.v3i1.17272

Bergougnoux, V. (2014). The history of tomato: From domestication to biopharming. In Biotechnology Advances (Vol. 32, Issue 1, pp. 170–189). Elsevier. 10.1016/j.biotechadv.2013.11.003

Chapagain, T., Khatri, B., & Mandal, J. L. (2011). Performance of Tomato Varieties during Rainy Season under Plastic House Conditions. Nepal Journal of Science and Technology, 12, 17–22. 10.3126/njst.v12i0.6473

Chaulagai, B., & Koirala, B. (2021). Review on major constraints during off seasonal tomato production in Nepal (Lycopersicon esculentum). International Journal of Horticulture and Food Science, 3(2). http://www.hortijournal.com

Coyne, D. L., Cortada, L., Dalzell, J. J., Claudius-Cole, A. O., Haukeland, S., Luambano, N., & Talwana, H. (2018). Plant-parasitic nematodes and food security in Sub-Saharan Africa. Annual Review of Phytopathology, 56(Volume 56, 2018, 381–403. 10.1146/ANNUREV-PHYTO-080417-045833/CITE/REFWORKS

de Araújo Filho, J. V., Machado, A. C. Z., Dallagnol, L. J., & Aranha Camargo, L. E. (2016). Root-knot nematodes (Meloidogyne spp.) parasitizing resistant tobacco cultivars in southern Brazil. Plant Disease, 100(6), 1222–1231. 10.1094/PDIS-03-15-0341-RE/ASSET/IMAGES/LARGE/PDIS-03-15-0341-RE_F4.JPEG

Devkota, S., Shrestha, S. L., Dhakal, D. D., Shakya, S. M., & Pandey, A. (2018). Evaluation of Tomato Hybrids for Yield Attributes Under Khumaltar Condiiton. Journal of Institute of Agriculture and Animal Sciences, 35, 191–196.

Eisenback, J., Hirschmann, H., & Triantaphyllou, A. C. (1980). Morphological Comparison of Meloidogyne Female Head Structures, Perineal Patterns, and Stylets. Journal of Nematology, 12(4), 300. /pmc/articles/PMC2618025/?report=abstract

El-Sappah, A. H., Islam, M. M., El-Awady, H. H., Yan, S., Qi, S., Liu, J., Cheng, G. T., & Liang, Y. (2019). Tomato Natural Resistance Genes in Controlling the Root-Knot Nematode. Genes, 10(925), 1–19. 10.3390/GENES10110925

FAO. (2011). The role of women in agriculture. ESA Working Paper No. 11-02. www.fao.org/economic/esa

*FAOSTAT*. (2021). http://www.fao.org/faostat/en/#data/QC

Ghimire, N., Kandel, M., Aryal, M., & Bhattarai, D. (2017). Assessment of Tomato Consumption and Demand in Nepal. The Journal of Agriculture and Environment, 18.

Ghimire, S., Subedi, P., & Green, S. (2001). Status of Tomato Yellow Leaf Curl Virus in Tomato in the Western Hills of Nepal. Nepal Agriculture Research Journal, 4 & *5*.

Hallmann, J., & Kiewnick, S. (2018). Virulence of Meloidogyne incognita populations and Meloidogyne enterolobii on resistant cucurbitaceous and solanaceous plant genotypes. Journal of Plant Diseases and Protection, 125, 415–424. 10.1007/s41348-018-0165-5

Hooks, C. R. R., Wang, K.-H., Ploeg, A., & Mcsorley, R. (2010). Using marigold (Tagetes spp.) as a cover crop to protect crops from plant-parasitic nematodes. Applied Soil Ecology, 46, 307–320. 10.1016/j.apsoil.2010.09.005

Javed, N., Gowen, S. R., El-Hassan, S. A., Inam-ul-Haq, M., Shahina, F., & Pembroke, B. (2008). Efficacy of neem (Azadirachta indica) formulations on biology of root-knot nematodes (Meloidogyne javanica) on tomato. Crop Protection, 27(1), 36–43. 10.1016/J.CROPRO.2007.04.006

Jones, J. T., Haegeman, A., Danchin, E. G. J., Gaur, H. S., Helder, J., Jones, M. G. K., Kikuchi, T., Manzanilla-López, R., Palomares-Rius, J. E., Wesemael, W. M. L., & Perry, R. N. (2013). Top 10 plant-parasitic nematodes in molecular plant pathology. Molecular Plant Pathology, 14(9), 946–961. 10.1111/MPP.12057

Khan, A., Ansari, S. A., Haris, M., Hussain, T., & Khan, A. A. (2023). Meloidogyne Species: Threat to Vegetable Produce. In Root-Galling Disease of Vegetable Plants (pp. 61–83). Springer, Singapore. 10.1007/978-981-99-3892-6_2

Khan, Z., Son, S. H., Akhtar, J., Gautam, N. K., & Kim, Y. H. (2012). Plant growth-promoting rhizobacterium (Paenibacillus polymyxa) induced systemic resistance in tomato (Lycopersicon esculentum) against root-knot nematode (Meloidogyne incognita). Indian Journal of Agricultural Sciences, 82(7), 603–610.

Luc, M., Sikora, R. A., & Bridge, J. (2005). Plant parasitic nematodes in subtropical and tropical agriculture: Second Edition. Plant Parasitic Nematodes in Subtropical and Tropical Agriculture: Second Edition, 1–871.

Lutuf, H., Nyaku, S. T., & Acheampong, M. (2018). Prevalence of plant-parasitic nematodes associated with tomatoes in three agro-ecological zones of Ghana. Ghana Journal of Agricultural Science, 52(1), 83–94. 10.4314/GJAS.V52I1

Magar, D. B. T., & Gauchan, D. (2016). Production, Marketing and Value Chain Mapping of “Srijana” Tomato Hybrid Seed in Nepal. Journal of Nepal Agricultural Research Council, 2, 1–8. 10.3126/jnarc.v2i0.16114

Manandhar, C., Baidya, S., Manandhar, S., Pant, B., Sharma, P., & Magar, P. (2020). Identification of Major Diseases of Tomato From Different Locations of Nepal. Journal of Plant Protection Society, 6.

Mandal, P., & Hossain, S. (2017). Status of agronomic strategies to control root-knot nematode (Meloidogyne spp.) in potato. Journal of Sylhet Agricultral University, 4(2), 161–172. https://www.researchgate.net/publication/333930280

MoALD. (2020). Statistical Information On Nepalese Agriculture 2075/76 (2018/19), Ministry of Agriculture & Livestock Development.

Ornat, C., & Sorribas, F. J. (2008). Integrated Management Of Root-Knot Nematodes In Mediterranean Horticultural Crops. In Integrated Management and Biocontrol of Vegetable and Grain Crops Nematodes (pp. 295–319). Springer, Dordrecht. 10.1007/978-1-4020-6063-2_14

Phadera, L. (2016). International Migration and its Effect on Labor Supply of the Left-Behind Household Members: Evidence from Nepal. Agricultural & Applied Economics Association.

Pokharel, R., Abawi, G. S., Zhang, N., Duxbury, J. M., & Smart, C. D. (2007). Characterization of Isolates of Meloidogyne from Rice-Wheat Production Fields in Nepal. Journal of Nematology, 39(3), 221. /pmc/articles/PMC2586498/

Pokharel, R., & Acharya, S. R. (2015). Sustainable Transport Development in Nepal: Challenges, Opportunities and Strategies. Journal of the Eastern Asia Society for Transportation Studies, 11, 209-.

Przybylska, A., & Obrępalska-Stęplowska, A. (2020). Plant defense responses in monocotyledonous and dicotyledonous host plants during root-knot nematode infection. Plant and Soil, 451, 239–260. 10.1007/S11104-020-04533-0/TABLES/2

Quinet, M., Angosto, T., Yuste-Lisbona, F. J., Blanchard-Gros, R., Bigot, S., Martinez, J. P., & Lutts, S. (2019). Tomato Fruit Development and Metabolism. In Frontiers in Plant Science (Vol. 10, p. 1554). Frontiers Media S.A. 10.3389/fpls.2019.01554

Seid, A., Fininsa, C., Mekete, T., Decraemer, W., & Wesemael, W. M. L. (2015). Tomato (Solanum lycopersicum) and root-knot nematodes (Meloidogyne spp.)-a century-old battle. Nematology, 17(9), 995–1009. 10.1163/15685411-00002935

Sikandar, A., Zhang, M. Y., Wang, Y. Y., Zhu, X. F., Liu, X. Y., Fan, H. Y., Xuan, Y. H., Chen, L. J., & Duan, Y. X. (2020). Review article: Meloidogyne incognita (root-knot nematode) a risk to agriculture. Applied Ecology and Environmental Research, 18(1), 1679–1690. 10.15666/aeer/1801_16791690

Sikora, R. A., Coyne, D., Hallman, J., & Timper, P. (Eds.). (2018). Plant Parasitic Nematodes in Subtropical and Tropical Agriculture, 3rd Edition (3rd ed.). CABI Publishing, Wallingford, UK. https://books.google.de/books?hl=en&lr=&id=mFloDwAAQBAJ&oi=fnd&pg=PR3&dq=%5BBOOK%5D+Plant+parasitic+nematodes+in+subtropical+and+tropical+agriculture+RA+Sikora,+D+Coyne,+J+Hallmann,+P+Timper+-+2018+&ots=cbB28e9UDy&sig=rYaCdJM_1Z3qh9L-iK7R2hnGakQ&redir_esc

Singh, K., Singh, A., & Singh, D. K. (1996). Molluscicidal activity of neem (Azadirachta indica A.Juss). Journal of Ethnopharmacology, 52(1), 35–40. 10.1016/0378-8741(96)01383-9

Speijer, P. R., & De Waele, D. (1997). Screening of Musa Germplasm for Resistance and Tolerance to Nematodes (1st ed.). INIBAP Technical Guidelines. https://books.google.de/books?hl=en&lr=&id=XAipDrKerBsC&oi=fnd&pg=PA3&dq=speijer+and+de+waele+1997&ots=jC1MAhTb0S&sig=Rs2dgS-QVZlPcq3Z9h1aZvHLdNw&redir_esc=y#v=onepage&q=speijeranddewaele1997&f=false

Taylor, D. P., & Netscher, C. (1974). An improved technique for preparing perineal patterns of Meloidogyne spp. Nematologica, 20(2), 268–269. 10.1163/187529274X00285

Thapaliya, K. P., Pyakuryal, K. N., Devkota, D., Panta, D., & Ghimire, A. (2023). Migration, Remittance and Its Consequences at the Household Level in Agrarian Nepal. International Journal of Agriculture Extension and Rural Development Studies, 10(2), 30–46. 10.37745/ijaerds.15/vol10n23046

Tiwari, I., Shah, K. K., Tripathi, S., Modi, B., Shrestha, J., Pandey, H. P., Bhattarai, B. P., & Rajbhandari, B. P. (2020). Post-harvest practices and loss assessment in tomato (Solanum lycopersicum L.) in Kathmandu, Nepal. Journal of Agriculture and Natural Resources, 3(2), 335–352. 10.3126/janr.v3i2.32545

Trudgill, D. L. (1997). Parthenogenetic root-knot nematodes (Meloidogyne spp.); how can these biotrophic endoparasites have such an enormous host range? Plant Pathology, 46(1), 26–32. 10.1046/J.1365-3059.1997.D01-201.X

Viaene, N., Coyne, D. L., & Davies, K. G. (2024). Biological and cultural management. In Plant Nematology (pp. 443–471). CABI International. 10.1079/9781800622456.0014

Viaene, N., Coyne, D. L., & Kerry, B. R. (2006). Biological and Cultural Management. Plant Nematology, 346–369.

Whipps, J. M., & Davies, K. G. (2000). Success in Biological Control of Plant Pathogens and Nematodes by Microorganisms. In G. Gurr & S. Wratten (Eds.), Biological Control: Measures of Success (pp. 231–269). Kluwer Academic Publishers. http://www.hri.ac.uk

Ye, W., Zeng, Y., & Kerns, J. (2015). Molecular Characterisation and Diagnosis of Root-Knot Nematodes (Meloidogyne spp.) from Turfgrasses in North Carolina, USA. PLOS ONE, 10(11), e0143556. 10.1371/JOURNAL.PONE.0143556

